## supplemental legends for "Morphogenetic forces planar polarize LGN/Pins in the embryonic head during *Drosophila* gastrulation"

**Figure 2- figure supplement 1. Pins is planar polarized on the posterior side of MD 3 and anterior side of MD 5 cells.** (A, C) Image of MD 3 (A) or MD 5 (C) of embryo expressing Pins::YFP, to label spindle rotation machinery, scalebar is 10 µm. (B, D) Quantification of average cellular (6-10 cells) Pins::YFP signal intensity around the mitotic cell cortex for 3 separate embryos, lines indicate average of cells in embryos for MD3 (B) or MD5 (D).

**Figure 3- figure supplement 1. Division angle is not correlated with interphase cell shape.** (A) Images are time series of MD1 of embryo with labeled Utr (Utr::GFP) magenta lines indicate division angle. Scalebar is 10 µm. **(B)** Divison angle is not correlated with interphase cells shape. Plot of the correlation between interphase cell shape and division angle., $R^2$=0.0002.

**Figure 3- figure supplement 2. Myristoylated Pins is uniformly recruited to the apical membrane.** (A, C) Image of MD1 of embryo expressing Pins::YFP, to label spindle rotation machinery (A), and Gap43::mCherry, to mark membranes (C). Scalebar is 10 µm. (B, D) Quantification of average cellular (10 cells) Pins::YFP (B) or Gap43::mCherry (D) signal intensity around the mitotic cell cortex for 3 separate embryos, lines indicate average of cells in all embryos for MD1. Cortical intensity was measured at anaphase, indicated by cell elongation, and normalized to cytoplasmic Pins::YFP (B) or Gap43::mCherry signal (D).

**Figure 3- figure supplement 3. Results from MDs 3 and 5 are consistent with results from MD1.** (A-D) Quantification of spindle angles in MDs 3 (A-B) and 5 (C-D) for Myristoylated Pins expressing embryos (B, D, UAS>myr::Pins::GFP) and control (A, C, UAS>Pins::GFP). Average orientation angles per embryo were statistically different between control and Myristoylated Pins expressing embryos for MDs 3 and 5, Mann-Whitney, p=0.0079 and 0.0317, respectively.

**Figure 4- figure supplement 1. Disruption of TLRs does not disrupt polarized division orientation** (A, C) Injection of TLR 2, 6, and 8 dsRNA does not disrupt division orientation. (A) Left. Image of MD1 of control injection (0.1x TE buffer injection) or (C) Image of MD1 of embryo injected with TLR 2, 6, and 8 dsRNA. Images show marker (Jup::GFP) to mark mitotic spindles (greyscale), scalebar is 10 µm. Right, same image annotated with dashed lines to indicate spindle position. (B, D) Quantification of spindle angles for control embryos (B, 0.1x TE buffer injection) or TLR 2, 6, 8 dsRNA injected embryos (D) using rose plots. For control embryos (0.1x TE buffer injection) MD1, n=8 embryos and 199 spindles. For TLR 2, 6, and 8 dsRNA injection MD1, n=4 embryos and 138 spindles. Average orientation angles per embryo were not statistically different between control and TLR 2, 6, and 8 dsRNA injection, Mann-Whitney, p=0.0635. Note 0.1x TE buffer is the same control and B the same plot as used for Figure 7B, but representative images are different. (E, F) Quantification of spindle angles for fixed control embryos (E, OR) and fixed TLR 2, 6, and 8 triple mutant embryos (F). For control embryos (OR) MD1, n=10 embryos and 246 spindles. For TLR 2, 6, and 8 triple mutant embryos MD1, n=3 embryos and 38 spindles. Average orientation angles per embryo were statistically different between control and TLR 2, 6, and 8 triple mutant embryos, Mann-Whitney, p=0.0070.

**Figure 4- figure supplement 2. Results from MDs 3 and 5 are consistent with results from MD1.** (A-D) Quantification of spindle angles in MDs 3 (A-B) and 5 (C-D) for $\alpha$–catenin depleted embryos (B, D, $\alpha$–catenin RNAi) and control (A, C, Rh3 RNAi). For $\alpha$–catenin depleted embryos MD3, n=6 embryos, 95 spindles, and for MD5, n=6 embryos, 129 spindles. For control (Rh3 RNAi) MD3, n=6 embryos, 120 spindles, and for MD5 n=6 embryos, 149 spindles. Average orientation angles per embryo were statistically different between control and $\alpha$–catenin depleted embryos for MDs 3 and 5, Mann-Whitney, p=0.0022 and 0.0043, respectively.

**Figure 6- figure supplement 1. Results from MDs 3 and 5 are consistent with results from MD1.** (A-D) Quantification of spindle angles in MDs 3 (A-B) and 5 (C-D) for CytoD injected embryos (B, D, 0.25mg/mL CytoD injected) and control (A, C, DMSO injection). For CytoD injected embryos MD3, n=6 embryos, 79 spindles, and for MD5, n=7 embryos, 101 spindles. For control (DMSO injection) MD3, n=5 embryos, 97 spindles, and for MD5 n=5 embryos, 90 spindles. Average orientation angles per embryo were statistically different between control and CytoD injected embryos for MDs 3 and 5, Mann-Whitney, p=0.0043 and 0.0025, respectively.

**Figure 6- figure supplement 2. Laser cutting schema and control.** (A, B) Schematics of laser ablation schema, AP cuts to disrupt forces from ventral furrow (A) or DV cut to disrupt forces from cephalic furrow (C). Separating MD1 from ventral furrow forces disrupts spindle orientation. Image of dorsal head of AP (B) or DV cut (D) embryos. Images show marker (Jupiter::GFP) to mark mitotic spindles (greyscale), scalebar is 50 µm. (E) For sham ablated embryos the ROI for was drawn around MD1 with laser power set to half ablation power. Sham ablated embryos were used as controls for ablation cut side, DV ablation, and AP ablation. For sham ablated embryos MD1, n-4 embryos and 58 spindles.
