## Supplementary figures and images for "Morphogenetic forces planar polarize LGN/Pins in the embryonic head during *Drosophila* gastrulation"

### Figure 2- figure supplement 1

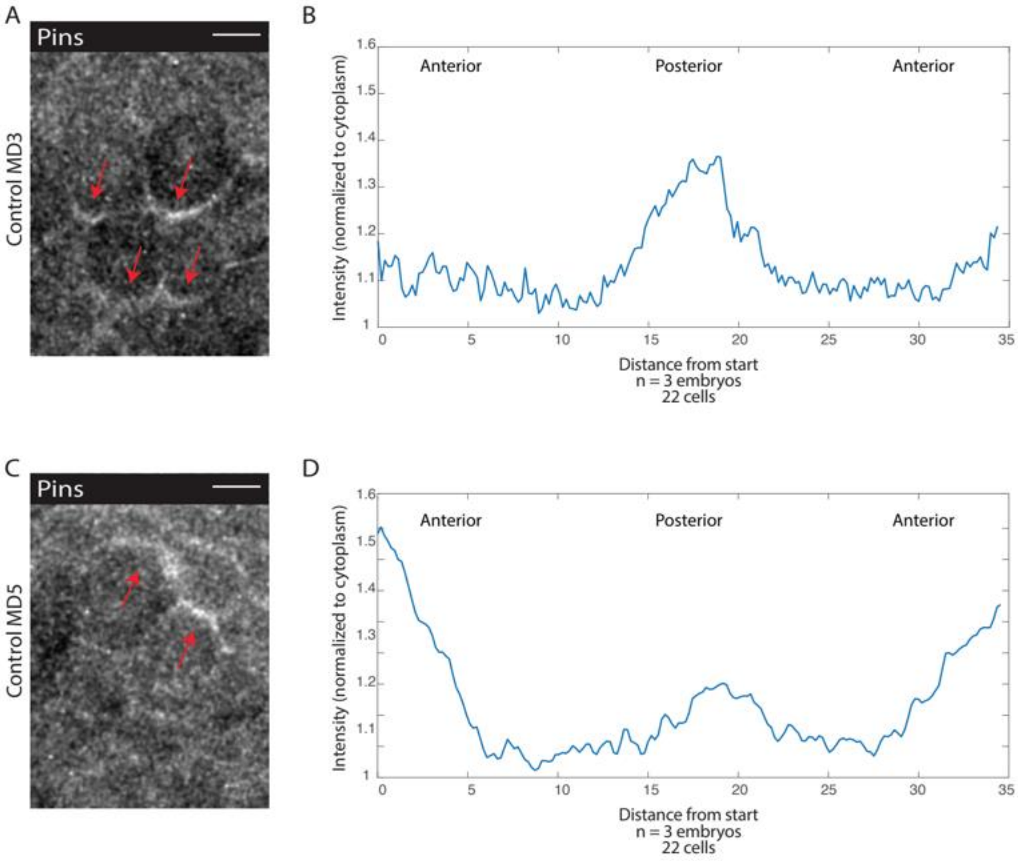

### Figure 3- figure supplement 1

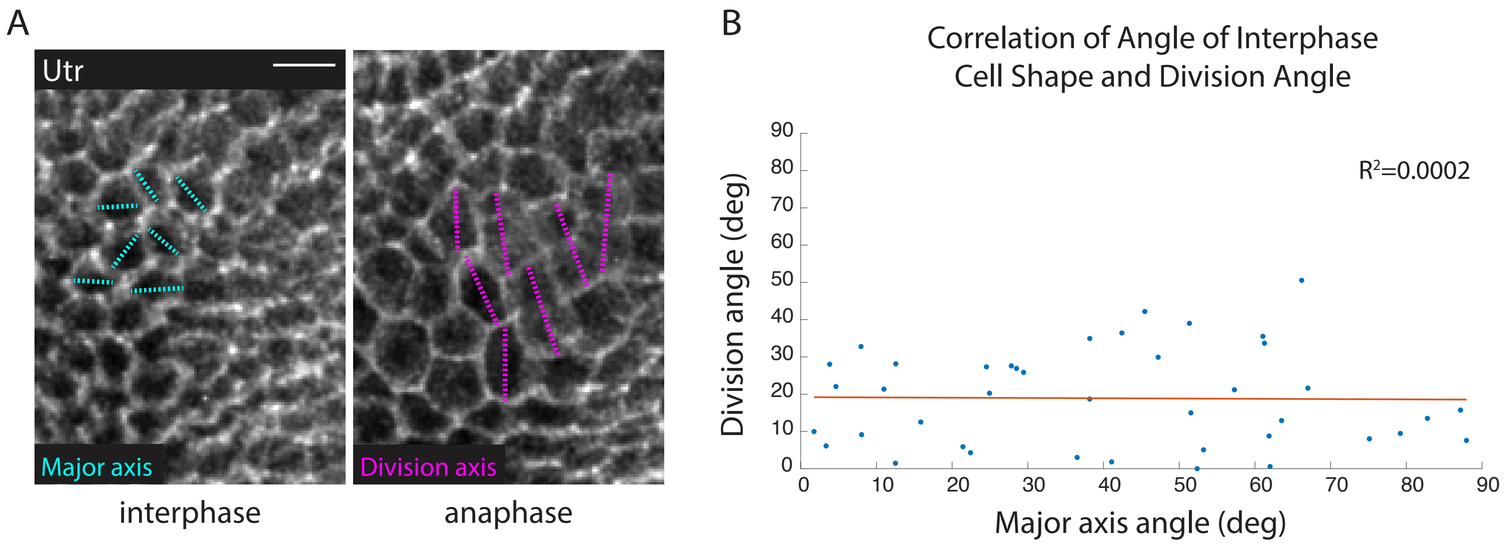

### Figure 3- figure supplement 2

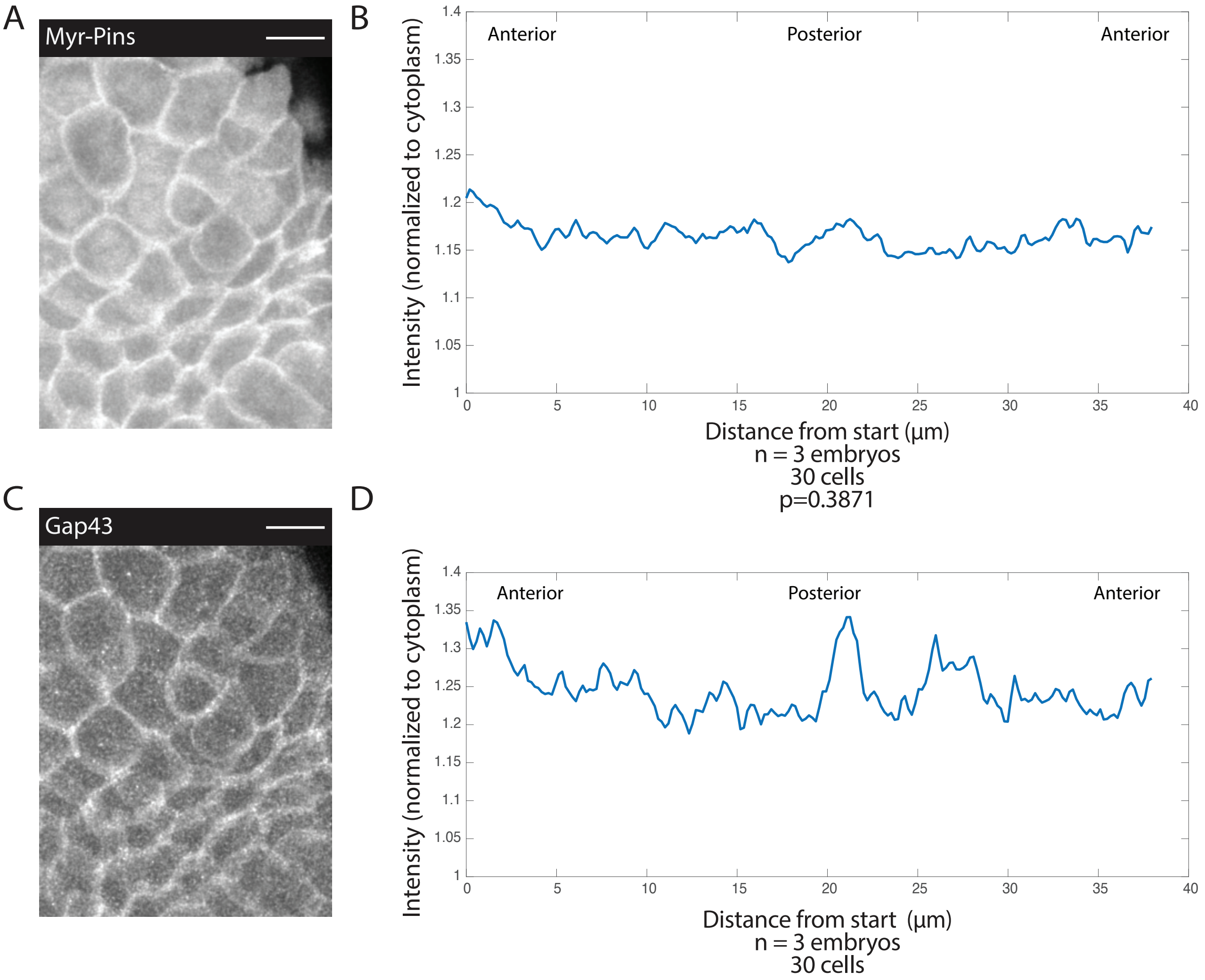

### Figure 3- figure supplement 3

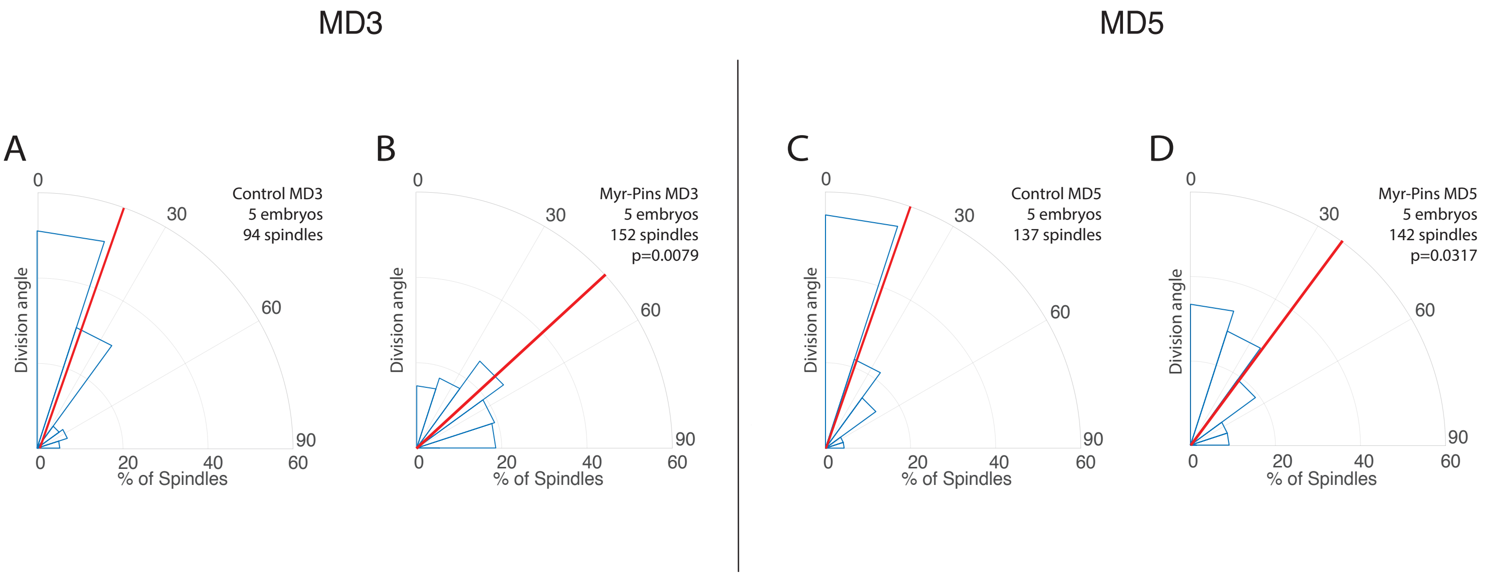

### Figure 4- figure supplement 1

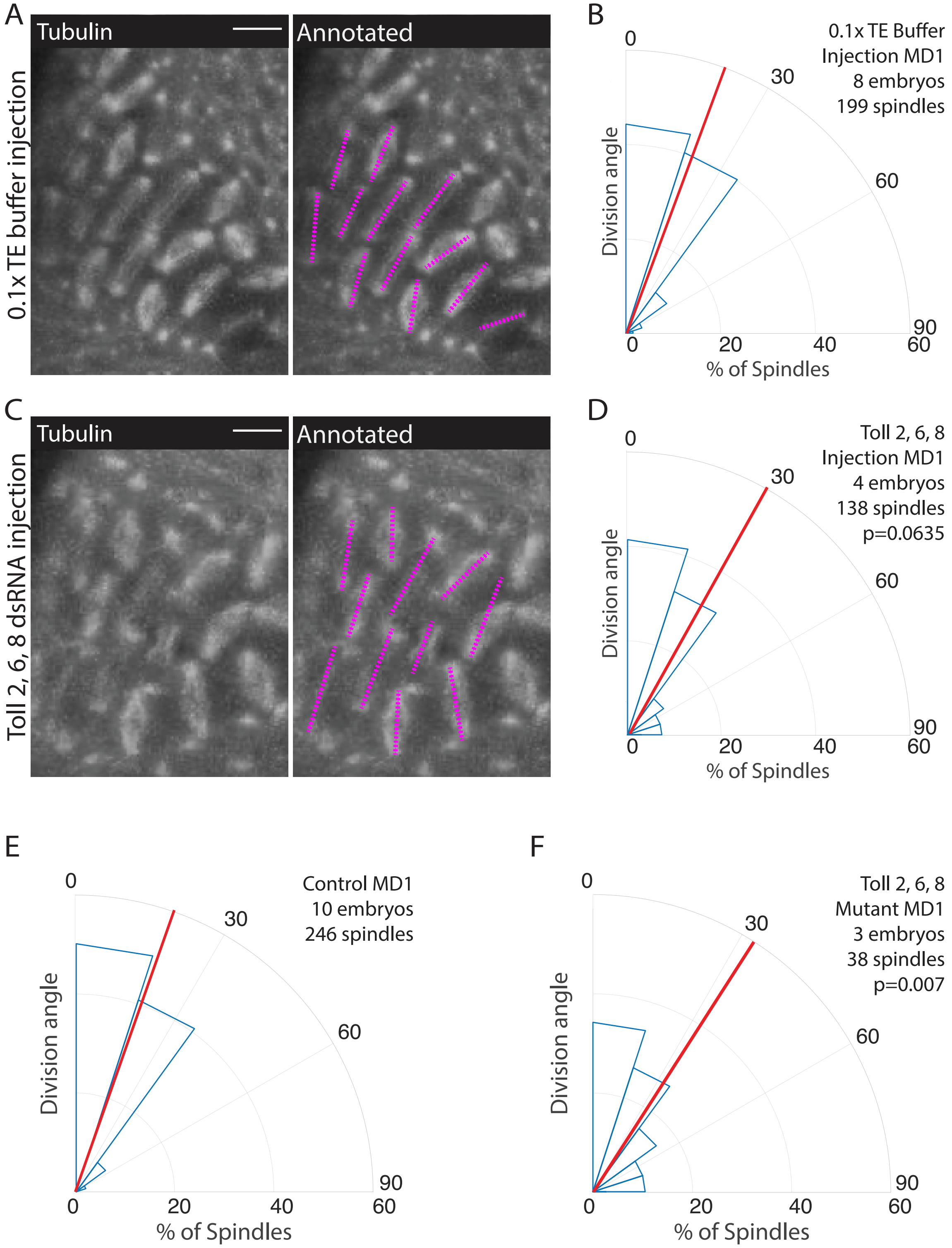

### Figure 4- figure supplement 2

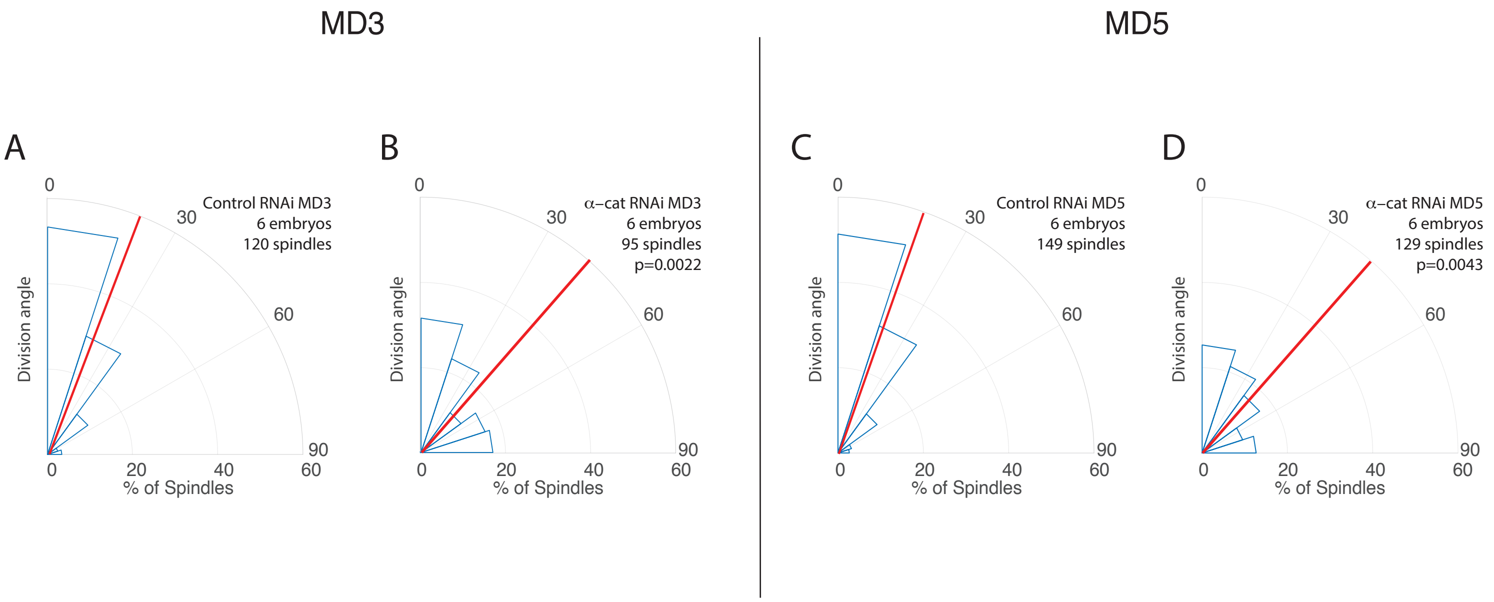

### Figure 6- figure supplement 1

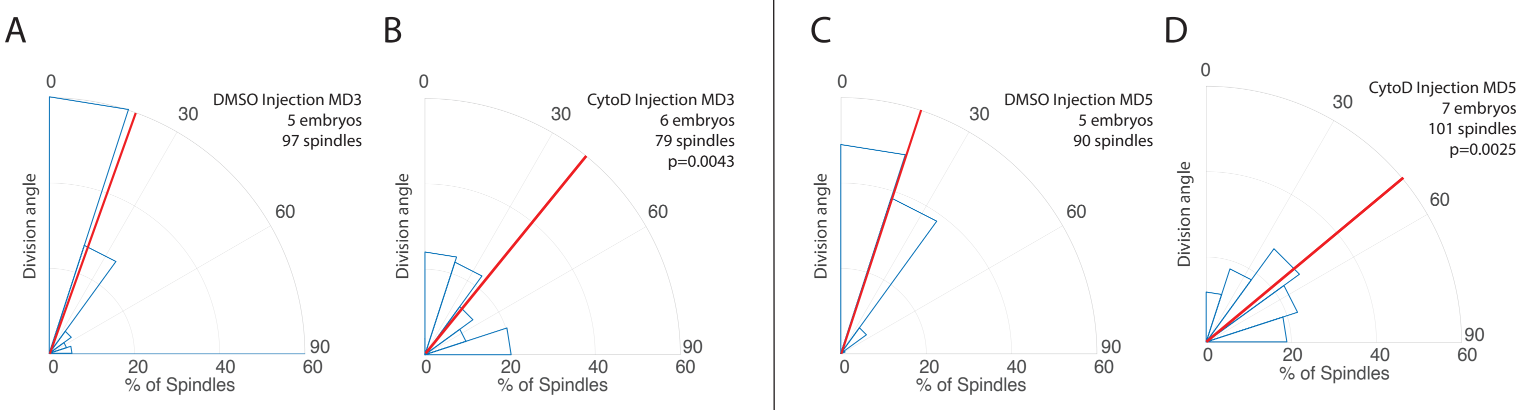

### Figure 6- figure supplement 2

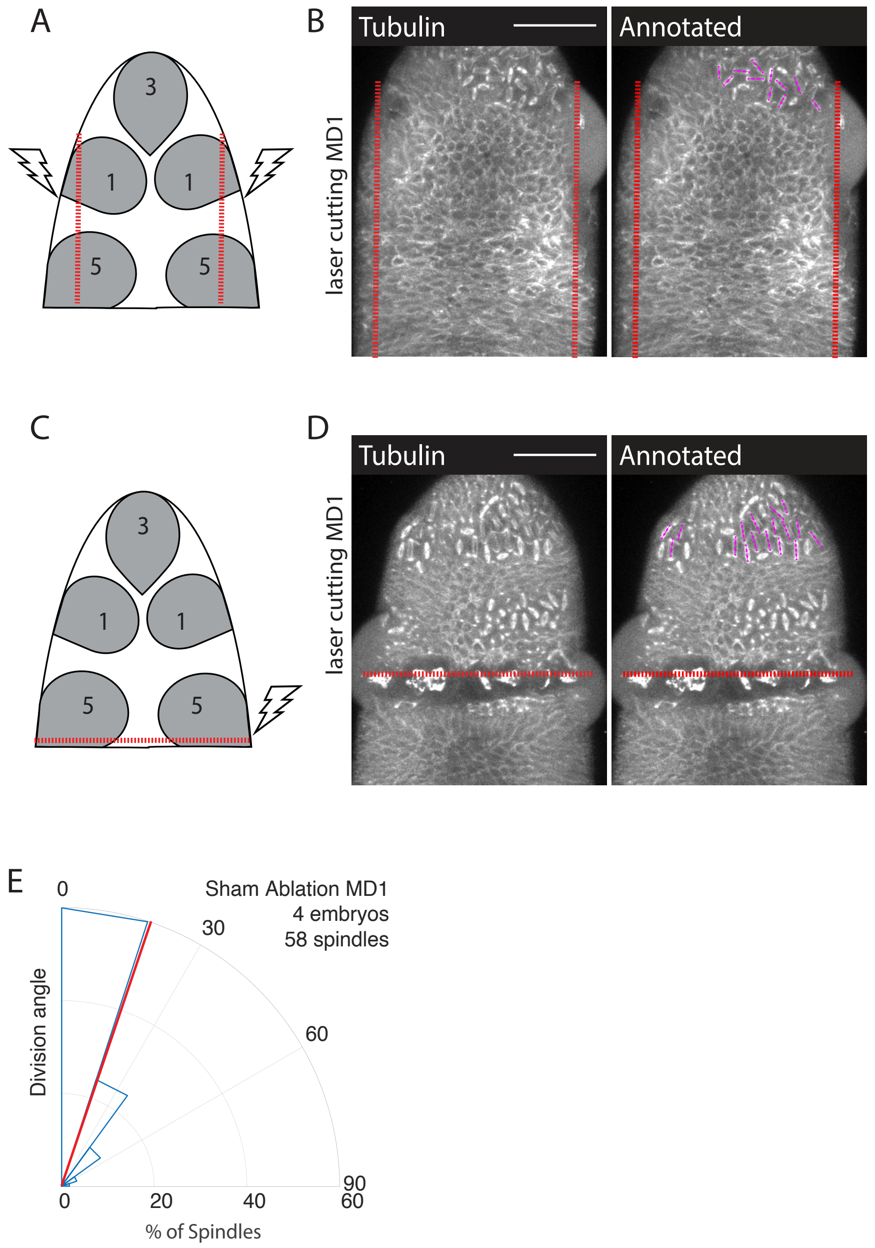
